# Synergy-informed bilevel optimization reveals joint-specific benefits for realistic exoskeleton torque prediction

**DOI:** 10.64898/2026.11.05.764547

**Authors:** Israel Luis, Thomas K. Uchida, Elena M. Gutierrez-Farewik

## Abstract

Simulation-driven optimization can guide exoskeleton control design, but its utility is limited by the accuracy with which muscle coordination is predicted. Strict effort-minimization formulations predict non-physiological muscle shutdown under high exoskeleton assistance, leading to peak assistive torques that exceed values reported in human-in-the-loop experiments. We evaluated the accuracy of synergy-informed simulations at predicting assistance across four single-joint exoskeleton configurations: ankle plantarflexion, knee extension, hip flexion, and hip abduction. We implemented a bilevel optimization to predict a parameterized assistive torque that minimizes metabolic cost, with assisted muscle dynamics estimated via either effort-minimization simulations (minimizing muscle effort only) or synergy-informed simulations (minimizing muscle effort constrained to a synergy structure). We used experimental data from five unimpaired adults walking at their preferred speed and compared results descriptively between the two formulations within each subject. Assistance based on effort-minimization simulations shut down agonist muscle activations and overestimated peak assistive torque compared with experimental values. In contrast, constraining muscle controls to a synergy structure prevented non-physiological muscle shutdown and yielded assistive torques closer to those measured in human-in-the-loop experiments. This benefit was not observed equally across all assisted degrees of freedom or subjects, emerging only where no single synergy could be suppressed to unload the assisted muscles. Synergy weights can therefore be interpreted as joint-specific predictors of when simulations can reliably inform wearable robot design.

**Author summary:** Exoskeletons are wearable robots that assist human movement. Personalized assistance can offer substantial benefits, but finding the best strategy requires testing many settings for each person, which is time-consuming. Computer simulations could speed up this search by identifying optimal assistance in a virtual environment. Yet simulations often predict much higher assistance than experiments find optimal because they tend to switch off assisted muscles, a strategy people do not adopt. One possible explanation is that simulations do not adequately represent how the nervous system coordinates muscles. Here, we examined whether simulating muscle control through synergies, groups of muscles activated together, produces more realistic predictions. Using walking data from five adults, we simulated assistance at the ankle, knee, and hip. Incorporating synergies kept key muscles active and brought predicted assistance closer to experimental values at the knee and, in three participants, the ankle, but not for hip flexion. Improvements depended on how the assisted muscles were coupled to other muscles within the synergies. These findings can help researchers assess when accounting for muscle coordination improves predictions used to guide exoskeleton design.

## Introduction

A central challenge in wearable robot design is predicting assistance strategies that will satisfy a given set of objectives. The potential benefits of exoskeletons have been demonstrated across a range of hardware configurations, from unpowered spring-based devices [1,2] to personalized motor-based devices that apply assistive torques across single joints [3] or multiple joints [4]. Despite these successes, the mechanisms underlying optimal assistive strategies remain difficult to interpret and predict. Experimental studies of various hardware–controller combinations have provided detailed descriptions of joint mechanics and muscle dynamics during assisted locomotion [5,6]. Yet, experimental studies typically focus on one or a small number of device configurations and operating conditions (e.g., walking on level ground at a controlled speed), making it time- and resource-intensive to establish general design principles. For systematic exploration of large hardware configuration or control design spaces, the experimental cost becomes intractable.

Musculoskeletal simulation offers a scalable alternative to physical prototyping. Prior work has demonstrated that simulation-based prediction of assistive torques can translate into measurable benefits in physical devices, including reduced metabolic cost [7], decreased knee joint loads [8], and improved balance under perturbations [9]. These results support the continued development of simulation-driven approaches as complements to experimental approaches. Nevertheless, a persistent and well-documented disagreement between predictions and experimental observations limits the reliability of simulation-based design. Effort-based formulations that minimize muscle activations are the most widely used approaches for simulating muscle coordination and human–robot adaptation. In an effort-based formulation, the cost function predicts muscle activations that are in good agreement with recorded electromyography (EMG) during unassisted walking [10]. However, it can predict physiologically implausible muscle coordination when simulating exoskeleton assistance. For example, when the objective is to minimize metabolic cost, the assistive torque leads to excessive deactivation of agonist or stabilizing muscles [11–16] even at the expense of recruiting antagonist muscles [11,12,16]. The optimization finds a mathematically valid but unrealistic solution: shut down muscles whose joint moments can be generated at lower metabolic cost. This is a coordination strategy that the human nervous system, which is subject to stability demands and sensorimotor constraints, is unlikely to adopt [17].

Muscle synergies provide a theoretical framework for modeling more realistic muscle coordination. According to this theory, the central nervous system coordinates movement through low-dimensional combinations of time-invariant synergy weights and time-varying synergy activations [18,19]. This modular organization has been proposed as a fundamental principle of muscle coordination that reduces the dimensionality of the motor command space and is well supported by electromyographic evidence during human locomotion [20–23]. In the context of wearable robotics, empirical evidence from robot-assisted gait has shown that wearers modulate the temporal activation of existing synergy modules rather than reorganizing their structure, indicating that synergy weight structures remain largely invariant when assistance is applied, particularly at moderate assistance levels [24–26]. This observation motivates modeling muscle coordination during assisted gait as a modulated form of synergy activations that are observed without assistance—that is, assuming an invariant synergy subspace rather than unconstrained muscle effort minimization. Importantly, during gait, effort minimization under biomechanical task constraints produces activation patterns that a small number of synergies can reconstruct and that closely match EMG-derived structures [27,28]. Therefore, synergy weights extracted from a walking simulation under an effort-minimization cost function can provide a physiologically representative constraint without requiring experimental EMG as input. This evidence motivates the examination of incorporating synergy modules into a framework based on minimal muscle effort when predicting optimal exoskeleton assistance.

In the context of gait, musculoskeletal simulations with synergy constraints have been shown to produce more realistic muscle recruitment patterns than unconstrained effort-based models [29–31]. However, to our knowledge, no previous study has isolated the effect of synergy constraints on the muscle coordination and optimal exoskeleton assistance predicted by musculoskeletal simulation, or compared the resulting assistive torques with those from human-in-the-loop experiments across multiple joints. Because synergy weights couple the activations of several muscles within a coordinated group, reducing the activation of one muscle necessarily modulates the activations of all the other muscles within the same synergy module, introducing additional constraints into the optimization problem that may prevent the shutdown of individual muscles.

Examining the isolated effect of synergy constraints on muscle coordination under assistance requires a simulation approach that fixes human–robot mechanics. Inverse simulation using experimental data collected during unassisted gait preserves joint kinematics and kinetics during the prediction of exoskeleton assistance; hence, this simulation approach provides an ideal testbed to investigate the effect of synergy constraints on effort minimization. By contrast, forward (predictive) simulation requires multi-objective criteria to replicate unassisted gait, which would obscure the analysis with confounding factors such as minimizing joint accelerations [32] or gait stability metrics [33].

The aims of this study were to evaluate whether constraining muscle control with synergy modules prevents muscle shutdown under optimal assistance prediction, and to examine whether a synergy-informed approach yields assistance torque profiles more consistent with those reported from human-in-the-loop experiments than a strictly effort-minimization approach. To this end, we implemented a synergy-informed muscle control model within an inverse simulation and embedded it within a bilevel optimization to predict joint-optimal exoskeleton assistance for four lower-limb degrees of freedom: ankle plantarflexion, knee extension, hip flexion, and hip abduction. We hypothesized that imposing synergy constraints would prevent the shutdown of individual muscles and would yield lower, more realistic peak assistive torques than those predicted using unconstrained effort minimization.

## Materials and methods

### Experimental data, musculoskeletal model, and joint mechanics

We used scaled musculoskeletal models, inverse kinematics solutions, and inverse dynamics solutions from a previous study [34], in which five unimpaired adults (2 male and 3 female, age: 31.4 ± 7.4 years, height: 1.75 ± 0.03 m, body mass: 69.0 ± 10.3 kg; mean ± standard deviation) walked at preferred speed (1.46 ± 0.07 m/s) along an instrumented walkway at the Promobilia MoveAbility Lab. Marker trajectories (Vicon V16, Oxford, UK) and ground reaction forces (AMTI Optima, Watertown, USA) were collected and processed in OpenSim version 4.4 [35]. The original study [34] was approved by the Swedish Ethical Review Authority (Dnr. 2020-02311) and all participants provided written informed consent. A scaled musculoskeletal model was created for each participant using a static calibration trial and the generic three-dimensional model by Rajagopal et al. [36], with adjustments proposed by Uhlrich et al. [37]; this model consisted of 29 degrees of freedom, including hip flexion/extension, hip abduction/adduction, hip internal/external rotation, knee flexion/extension, and ankle plantarflexion/dorsiflexion, and 40 lower-limb muscles per leg (Fig 1A). Joint kinematics and kinetics were computed using OpenSim’s Inverse Kinematics and Inverse Dynamics tools.

**Fig 1.**
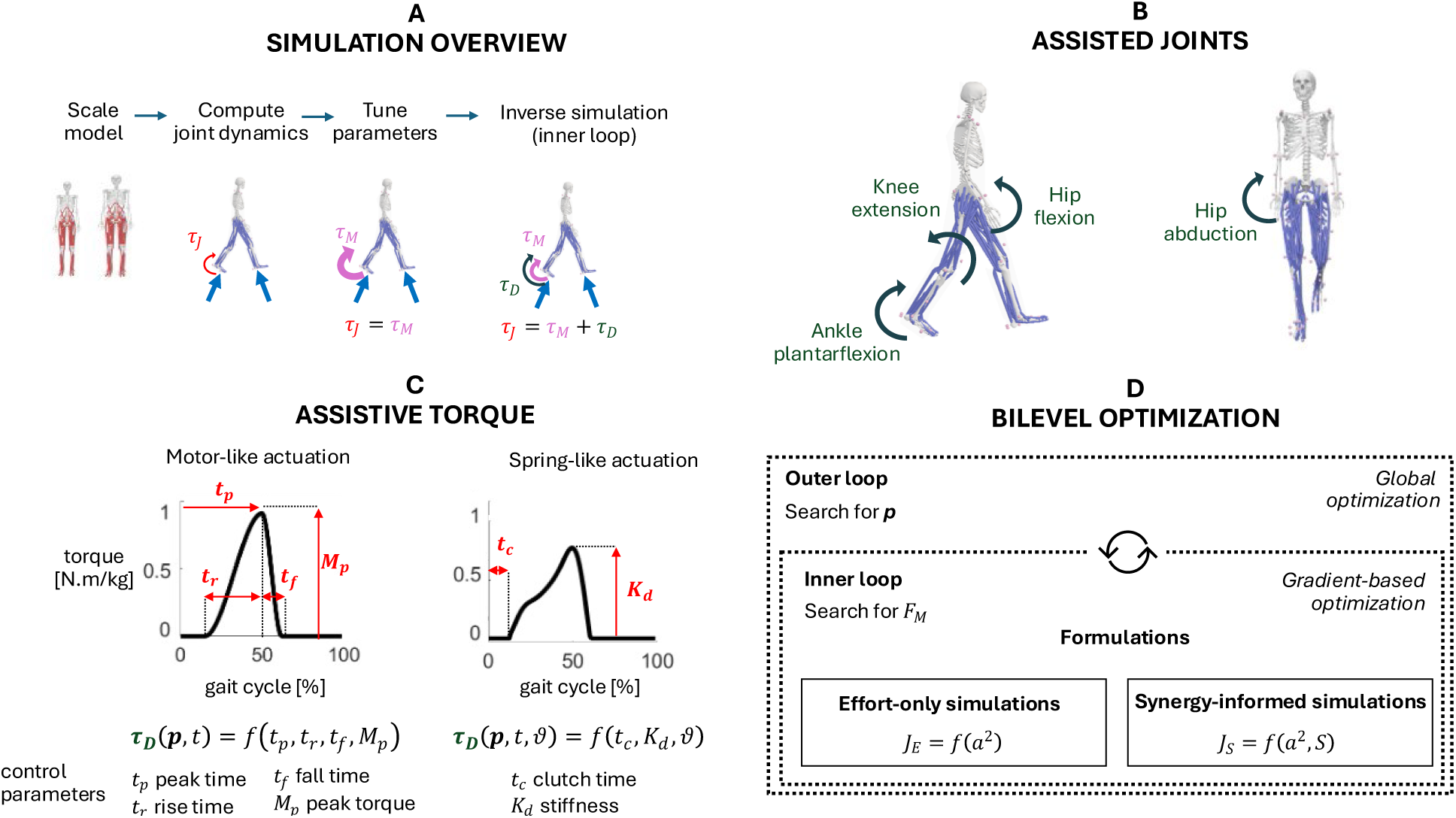
Summary of methods. (A) Simulation overview: A generic musculoskeletal model was scaled based on a calibration trial. Inverse kinematics and inverse dynamics were computed using experimental motion data. Musculotendon parameters were tuned based on experimental data reported in the literature. The tuned model and prescribed joint mechanics were used in the inner loop of the bilevel optimization. (B) Assisted joints: Ankle plantarflexion, knee extension, hip flexion, or hip abduction. (C) Assistive torque: Motor-like actuation was defined by 4 parameters describing a torque–time profile; spring-like actuation, by 2 parameters describing a torque applied based on the joint angle (*θθ*) computed from inverse kinematics. (D) Bilevel optimization: Nested optimizations where the inner loop searched for muscle forces (*F_M_*) that solved the muscle redundancy problem, and the outer loop searched for control parameters (***p***) that minimized the average metabolic cost during gait. Two formulations of the inner loop were evaluated: effort-only simulations, which minimized the sum of squared muscle activations (*a*^2^), and synergy-informed simulations, which additionally encouraged adherence of muscle activations to synergy modules (*S*).

Musculotendon parameters were adjusted using a two-phase optimization method [38] to resemble reported experimental observations. Optimal fiber lengths, tendon slack lengths, and tendon stiffnesses of the gastrocnemius, soleus, and vastus muscles were optimized to minimize the difference between simulated fiber lengths and those reported from ultrasonography [39,40]. Parameters defining the passive force–length relationships were optimized to minimize the difference between simulated passive moments and *in vivo* data [41]. This population-derived calibration method provided closer agreement between simulated muscle excitations and EMG recordings during gait than linearly scaled musculotendon parameters [38].

### Assistive devices

We simulated four single-joint devices, each assisting one degree of freedom in one direction: ankle plantarflexion, knee extension, hip flexion, and hip abduction (Fig 1B). All devices were modeled as ideal torque actuators. Simulated devices providing ankle plantarflexion, hip flexion, and hip abduction assistance generated motor-like torque profiles parameterized by peak time (*t_p_*), rise time (*t_r_*), fall time (*t_f_*), and peak magnitude (*M_p_*); the knee extension device provided a spring-like torque profile parameterized by clutch time (*t_c_*) and rotational stiffness (*K_d_*) (Fig 1C). These parameterizations were used in human-in-the-loop optimization (HILO) studies [4,42,43] and enabled direct comparison with previously reported optimal torque profiles.

Actuators were modeled as ideal torque generators to isolate simulated muscle coordination from implementation-specific factors such as actuator mass, bandwidth limitations, and efficiency losses. Actuator torque profiles *τ_D_* were functions of the control parameter vector ***p***. Each optimal control vector ***p*^∗^** was determined via bilevel optimization (Fig 1D), as described in the Bilevel optimization section.

### Musculoskeletal simulation

We simulated muscle dynamics under torque assistance using two formulations: effort-only simulation and synergy-informed simulation. Both formulations were implemented using inverse simulation with direct collocation optimal control, based on an algorithm proposed by De Groote et al. [44]. Joint kinematics and kinetics of the musculoskeletal simulation were prescribed using the previously computed inverse kinematics and inverse dynamics solutions, and muscle dynamics were simulated by solving an optimization problem, described in detail for each formulation in the following two subsections.

### Effort-only simulation

This formulation simulated muscle dynamics without imposing a synergy structure, and it has been adopted in previous studies [12,44]. In brief, the state variables were muscle activations *a*(*t*) and normalized fiber lengths *IỸ_M_*(*t*). Controls were muscle excitations *e*(*t*) and auxiliary controls were normalized fiber velocities *vỸ_M_*(*t*). Cost function *J_E_* was a weighted multi-objective cost function that primarily minimized the sum of squared muscle activations *C_E_*. The cost function included two additional terms: *C_R_*, which penalized the use of reserve actuator *τ_R_* (a non-physiological torque generated when muscle–tendon actuators cannot produce the torques computed from inverse dynamics), and *C_V_*, which provided numerical regularization:

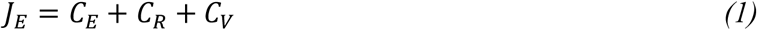

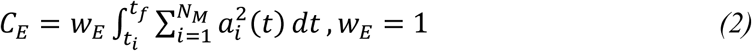

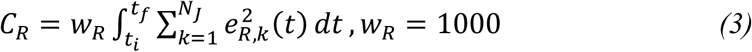

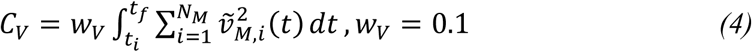

where *t_i_* and *t_f_* were the initial and final times of the simulation, *N_M_* was the number of muscles per leg (*N_M_* = 40), *N_J_* was the number of degrees of freedom per leg (*N_J_* = 5), and *e_R_*_,*k*_(*t*) was the reserve actuator excitation at degrees of freedom *k* and was bounded between -1 and +1. Weights *w_E_*, *w_R_*, and *w_V_* were selected to balance the three objectives *C_E_*, *C_R_*, and *C_V_* [44]. Equilibrium was imposed at each degree of freedom and at each time point between the joint torque *τ_J_* from inverse dynamics and the sum of muscle-generated moments *τ_M_*, reserve actuator torques *τ_R_*, and assistive torque *τ_D_* (either *τ_D_*(***p***, *t*) or *τ_D_*(***p***, *t*, *θ*) where applicable):

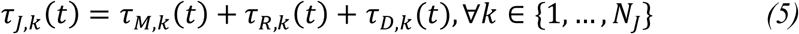

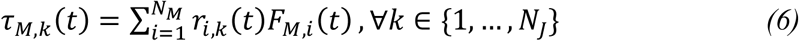

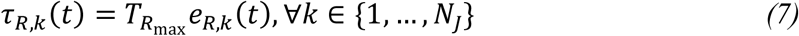

where *r_i_*_,*k*_ was the moment arm of the muscle *i* about the degree of freedom *k*, *F_M_*_,*i*_ was the force generated by the muscle *i*, *T_R_*_max_ was the maximum reserve actuator torque (*T_R_*_max_ = 100 N⋅m), and *τ_D_*_,*k*_ was the assistive torque, which was nonzero only at the assisted degree of freedom *k*. Additionally, activation dynamics and contraction dynamics were imposed as constraints, and an equilibrium was imposed between tendon and muscle forces. A detailed description of this formulation can be found in De Groote et al. [44].

### Synergy-informed simulation

We implemented a simulation algorithm with synergy-informed muscle control by extending the effort-only formulation described above, constraining muscle activations to adhere to a synergy structure. We constrained muscle activations rather than muscle excitations for a computational reason: muscle activation is a first-order function of muscle excitation, so activation attenuates the abrupt changes that excitation can exhibit at the direct collocation mesh points; constraining the smoother variable (activation) improved the numerical conditioning of the optimization problem.

We computed muscle dynamics in the unassisted condition by using the effort-only simulation with *τ_D_* = 0 (Eq 5). The result was a “baseline” muscle activation matrix *a_U_* ∈ R^(*N*^*^M^*^×*T*)^, representing the nominal, minimal-effort coordination pattern. Then, muscle synergies were extracted by applying non-negative matrix factorization (NMF) to the baseline activation matrix *a_U_*, yielding the decomposition *a_U_*(*t*) ≈ ***W**_U_**H**_U_*(*t*), where ***W**_U_* ∈ R^(*N*^*^M^*^×*N*^*^S^*^)^ was the matrix of time-invariant synergy weights, ***H**_U_*(*t*) ∈ R^(*N*^*^S^*^×*T*)^ was the matrix of time-varying synergy activations, and *N_S_* was the number of synergies. NMF was performed using an alternating least squares strategy with 100 random initializations and 2000 iterations per initialization to ensure robust factorization and avoid convergence to local minima. The extraction required specifying the number of synergies, which is described in the Data analysis section.

A fundamental modeling choice was that the synergy weights ***W**_U_* were invariant between unassisted and assisted walking conditions, consistent with empirical evidence from robot-assisted gait [24–26]. Under this assumption, changes in muscle activations during assistance were modulated by synergy activations, computed as *a_A_*(*t*) = ***W****_U_**H**_A_*(*t*), where ***H**_A_*(*t*) were the adapted synergy activations. We implemented this muscle coordination strategy by simulating muscle dynamics as in the effort-only simulations and including a synergy deviation penalty in the objective function:

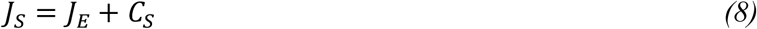

where *C_S_* was a term that encouraged muscle activations to adhere to the subspace defined by ***W****_U_*:

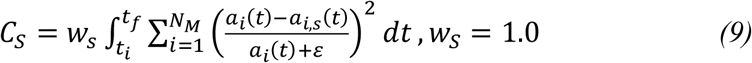

The projected activation *a_i_*_,*s*_(*t*) = [***W****_U_**H**_A_*(*t*)]*_i_* was computed from the invariant synergy weight ***W****_U_*, which was defined as in the unassisted condition, and the adapted synergy activation ***H**_A_*(*t*), which was a free variable in the optimization. The denominator normalized the penalty across muscles; *ε* = 0.01 was added for numerical stability. The weight *w_S_* = 1.0 was selected to encourage adherence to the synergy modules while permitting some modulation of synergy activations under assistance. This formulation penalized relative rather than absolute deviation from the synergy subspace, ensuring that the cost of deviating from the synergy projection remained proportional to the local activation magnitude. A qualitative comparison of our approach with existing formulations [31,45–47], as well as the analysis used to select *w_S_*, is provided in S1 Appendix (section A). The algorithms for the effort-only and synergy-informed simulations were implemented in MATLAB R2024a using a customized version of the direct collocation framework by De Groote et al. [44].

### Bilevel optimization

The computation of optimal assistive torque was formalized as a bilevel optimization problem (Fig 1D). The inner loop searched for muscle controls that minimized the cost function used to solve the muscle redundancy problem. This optimization provided simulated muscle dynamics for a given assistive torque *τ_D_*, as well as the “physiological” matrix ***χ***^∗^(***p***) containing the time histories of all simulated states, state derivatives, and muscle controls. The outer loop searched for control parameters *p* that minimized the cost function, which was a function of the assistive goal (computed from ***χ***^∗^(***p***)).

We constructed one bilevel optimization per simulation formulation (i.e., effort-only and synergy-informed) and accounted for stride-to-stride variability in gait [48]. It is common practice to predict optimal assistance for several (e.g., 3 to 5) gait cycles individually and then average the results to obtain a representative solution [11–15]. However, this approach poses a methodological challenge: a torque profile that has been averaged over several gait cycles will not be optimal for any individual gait cycle and may obfuscate comparisons. Therefore, we evaluated the muscle dynamics of a single assistive torque profile applied identically across three gait cycles, each with its own joint kinematics and kinetics. This multi-cycle approach promoted an assistive torque representative across multiple gait cycles, accounting for stride-to-stride variability during steady-state walking and is consistent with conditions used in many human-in-the-loop optimization studies.

For each candidate torque profile, the inner loop simulated muscle dynamics across three gait cycles using either the effort-only or the synergy-informed formulation, and the outer loop used the resulting physiological matrices to compute a per-stride assistive goal, which was then averaged to obtain a representative solution. This multi-cycle representation was particularly important for the synergy-informed simulations, as synergy activations were free to vary independently across gait cycles in response to the same assistive torque, while the synergy weights remained constant. Our approach reflected the ability of the central nervous system to modulate cycle-to-cycle coordination within an invariant high-order synergy module.

The assistive goal was the time-averaged whole-body metabolic cost, which was computed from physiological matrices and a metabolic energy model from the literature. Specifically, we computed the instantaneous metabolic energy rate (i.e., muscle metabolic power *Ė_M_*(*t*)) for each muscle in the musculoskeletal model using the model of Bhargava et al. [49]. We assumed that negative muscle power did not contribute to the metabolic power, such that metabolic power was non-negative at all instants of time. We computed the total metabolic power in one leg *Ė_M_*(*t*) as the sum of all muscles’ metabolic power:

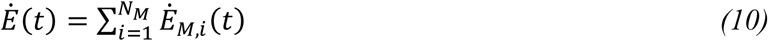

The average metabolic cost *E̅* was computed as the integral of the total metabolic power in one leg *Ė*(*t*) divided by the gait cycle duration and doubled, assuming the right and left legs incurred the same metabolic cost:

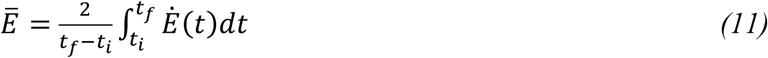

The outer loop was implemented using Bayesian optimization (MATLAB bayesopt, Statistics and Machine Learning Toolbox). Initial guesses were generated using a stratified strategy to span a diverse, physically feasible parameter space. For motor-like actuation, initial guesses were stratified over the peak torque *M_p_* and peak time *t_p_* (Fig A in S2 Appendix). The feasible ranges of *M_p_* and *t_p_* were each divided into three intervals, forming a 3×3 grid, and each of the 9 cells was populated with 3 four-parameter vectors (peak time, rise time, fall time, and peak torque), yielding 27 initial guesses. A batch of candidate vectors was drawn by Latin Hypercube Sampling over the full four-parameter space, sorted into cells according to which *M_p_* and *t_p_* interval each vector fell into, and retained only if they satisfied temporal feasibility constraints; batches were redrawn until each cell held at least three feasible candidates. Following this process, within each cell, the three vector candidates with the most evenly spread *M_p_* values were retained as the initial guesses. Temporal feasibility constraints, enforced as linear inequality constraints, limited assistive torques to be applied within certain gait phases, as in human-in-the-loop experiments [4,42,43]. For spring-like actuation, we generated 12 initial guesses via Latin Hypercube Sampling over the full two-parameter space (clutch time and stiffness). The maximum number of evaluations was set to 50 iterations per control parameter, yielding 200 iterations for motor-like actuation (four parameters) and 100 iterations for spring-like actuation (two parameters). The parameter bounds for each actuator type and temporal feasibility constraints for motor-like assistance are provided in S1 Appendix (section B). Simulations were run in MATLAB R2024a on an AMD Ryzen 7 7800X3D processor (8 physical cores, 4.20 GHz) with 32 GB RAM. Computation was parallelized across 8 workers via the Parallel Computing Toolbox.

### Data analysis

We first evaluated the synergy-informed simulation without assistance to establish the physiological plausibility of the predicted muscle activations prior to introducing assistance. For each participant, we tested three synergy dimensions (*N_S_*): 4, 5, and 6, which spanned the range of synergy counts reported for human walking [45]. The number of synergies was selected as the minimum required to achieve a variance accounted for (VAF) of at least 95% relative to the minimal-effort baseline, averaged across all participants. We reported two synergy reconstruction quality metrics (VAF and root mean square error, RMSE), muscle activation, and average metabolic cost across all synergy dimensions to characterize the sensitivity of the simulation to the number of synergies. We compared the predicted muscle activations with EMG reported by Perry and Burnfield [50] and the predicted average metabolic cost with indirect calorimetry collected in a previous study [34]. We then continued the analysis with the synergy dimension that met the reconstruction criterion.

We reported synergy weights and activations when unassisted to characterize the inter-subject consistency of the extracted modules and their association with major muscle groups. Because NMF does not guarantee a fixed ordering of the extracted modules, we reordered synergies across participants using the Hungarian algorithm [51], applied to the cosine similarity matrix of synergy weights, to enable qualitative comparison.

We then used bilevel optimization to compute optimal assistance for each of the four targeted degrees of freedom: ankle plantarflexion, knee extension, hip flexion, and hip abduction. For each assistive device, we reported and compared the predicted optimal torque profiles and control parameters against values reported from experiments [4,43,52,53], and presented the predicted reductions in average metabolic cost relative to unassisted conditions corresponding to each subject. To characterize the differences in muscle dynamics predicted by the two formulations, we presented activations of the primary agonist and antagonist muscles and synergy activations at each assisted joint. No inferential statistical tests were performed; given the sample of five subjects, results were described for each subject and were not tested for statistical significance.

## Results

### Muscle activations and average metabolic cost between formulations

Increasing the synergy dimension improved the reconstruction of the muscle activations in the synergy-informed simulations. The VAF increased from 90% with 4 synergies (RMSE = 0.029) to 95% with 6 synergies (RMSE = 0.021; Table 1). Qualitatively, the timing of muscle activations was similar between the two formulations, even across different synergy dimensions, and compared favorably with EMG timing (Fig B in S2 Appendix). In the synergy-informed simulations, the magnitudes of activations and co-activations decreased as the synergy dimension increased, as would be expected by increasing the dimensionality of the control space, and the tendency was more pronounced in certain muscles (Fig B in S2 Appendix).

**Table 1.**
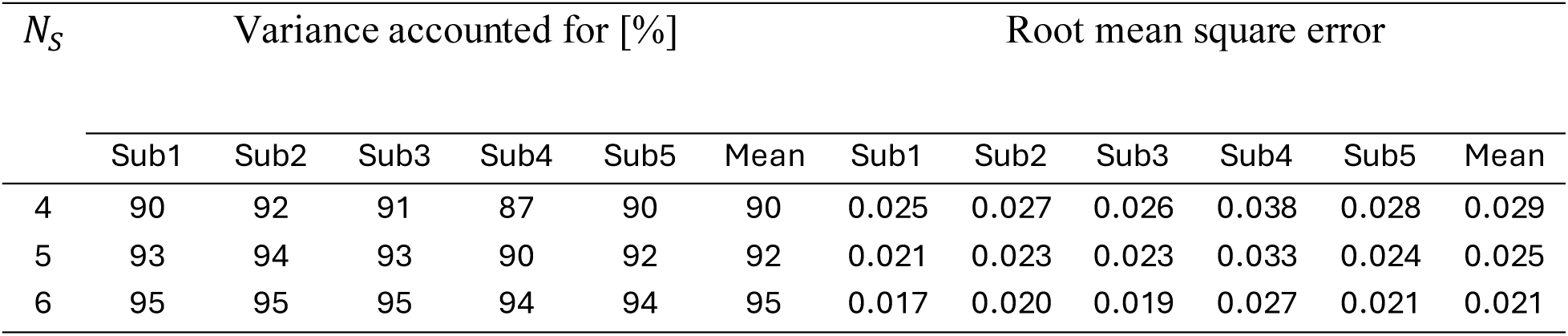
Synergy reconstruction quality across synergy dimensions. Subject-specific and mean values across five subjects (“Sub1” to “Sub5”). *N_S_* : number of synergies.

| *N*<sub>*S*</sub> | Variance accounted for [%] |  |  |  |  |  | Root mean square error |  |  |  |  |  |
|  | Sub1 | Sub2 | Sub3 | Sub4 | Sub5 | Mean | Sub1 | Sub2 | Sub3 | Sub4 | Sub5 | Mean |
|---|---|---|---|---|---|---|---|---|---|---|---|---|
| 4 | 90 | 92 | 91 | 87 | 90 | 90 | 0.025 | 0.027 | 0.026 | 0.038 | 0.028 | 0.029 |
| 5 | 93 | 94 | 93 | 90 | 92 | 92 | 0.021 | 0.023 | 0.023 | 0.033 | 0.024 | 0.025 |
| 6 | 95 | 95 | 95 | 94 | 94 | 95 | 0.017 | 0.020 | 0.019 | 0.027 | 0.021 | 0.021 |

The predicted average metabolic cost decreased as the synergy dimension increased. Effort-only simulations predicted an average metabolic cost of 5.3 W/kg (normalized by body mass), which was 13% greater than experimental measurements (4.7 W/kg; Fig B in S2 Appendix). Relative to experimental data, the predicted average metabolic cost was 38% greater with 4 synergies (i.e., 6.5 W/kg) and 32% greater with 6 synergies (i.e., 6.2 W/kg). Relative to effort-only simulations, the overestimation ranged from 17% to 23% as the synergy dimension increased from 4 to 6.

The simulation with 6 synergies met the reconstruction criterion. This synergy dimension provided high reconstruction accuracy and good temporal agreement with experimental EMG, primarily in muscles crossing the ankle and knee joints, yet an average metabolic cost prediction greater than that observed experimentally. We used effort-only simulations and synergy-informed simulations based on 6 synergies for all subsequent analyses.

### Metabolic cost profiles between formulations

Average metabolic costs predicted by the synergy-informed simulations were greater than those of the effort-only simulations, yet the timing of the metabolic power peak was similar between formulations. Both formulations predicted three metabolic power peaks within the gait cycle, yet synergy-informed simulations predicted a consistently greater magnitude throughout the gait cycle (Fig 2). The two formulations predicted similar muscle contributions: the soleus dominated metabolic cost among the ankle plantarflexors; the vastus lateralis, among the knee extensors; the psoas and rectus femoris, among the hip flexors; and the gluteus medius, among the hip abductors (Fig 3).

**Fig 2.**
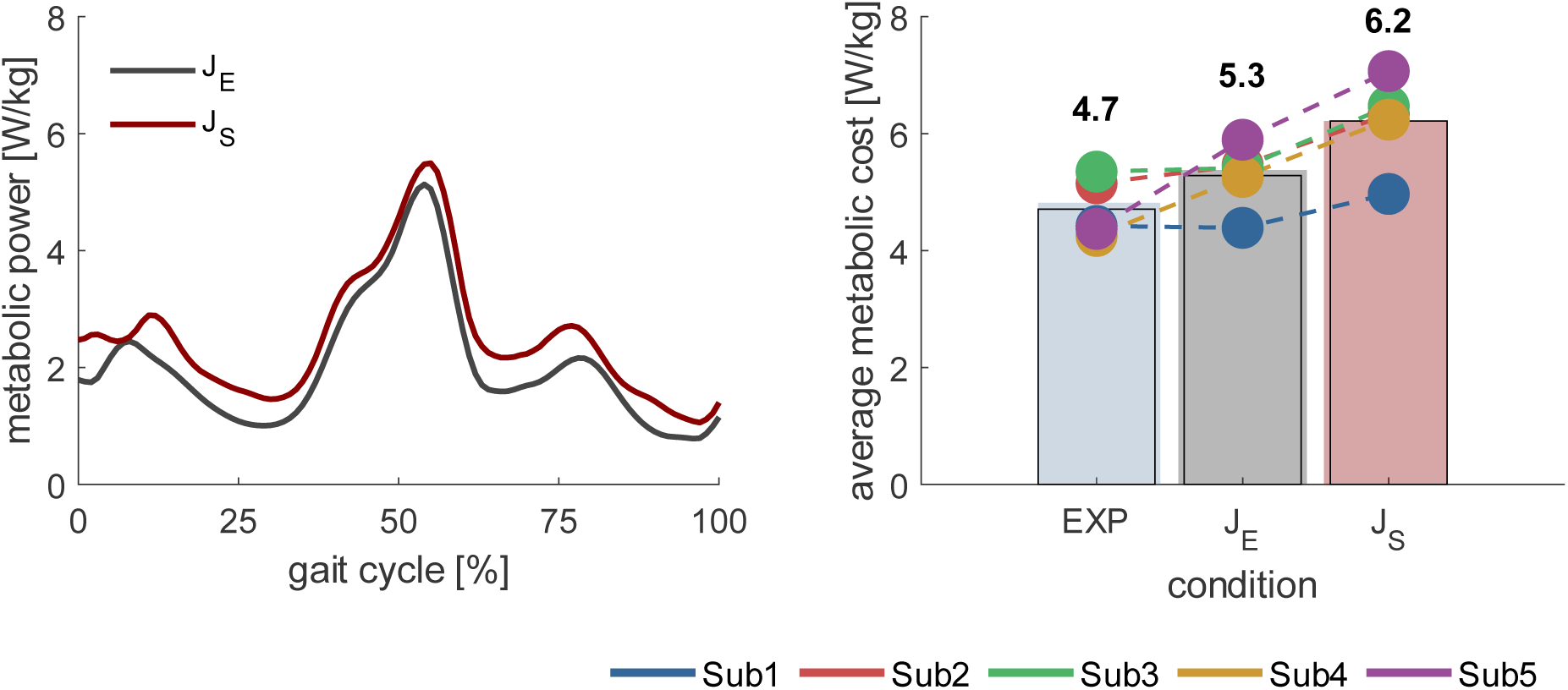
Metabolic power and average metabolic cost in unassisted simulations. Metabolic power in one leg (left) and average metabolic cost (right) were predicted by effort-only simulation (J_E_) and synergy-informed simulation (J_S_). Experimental metabolic cost (EXP) was measured using indirect calorimetry [34] and is shown for comparison. Mean values were computed across all subjects and gait cycles. Gait cycle was defined from heel strike to heel strike of the same limb.

**Fig 3.**
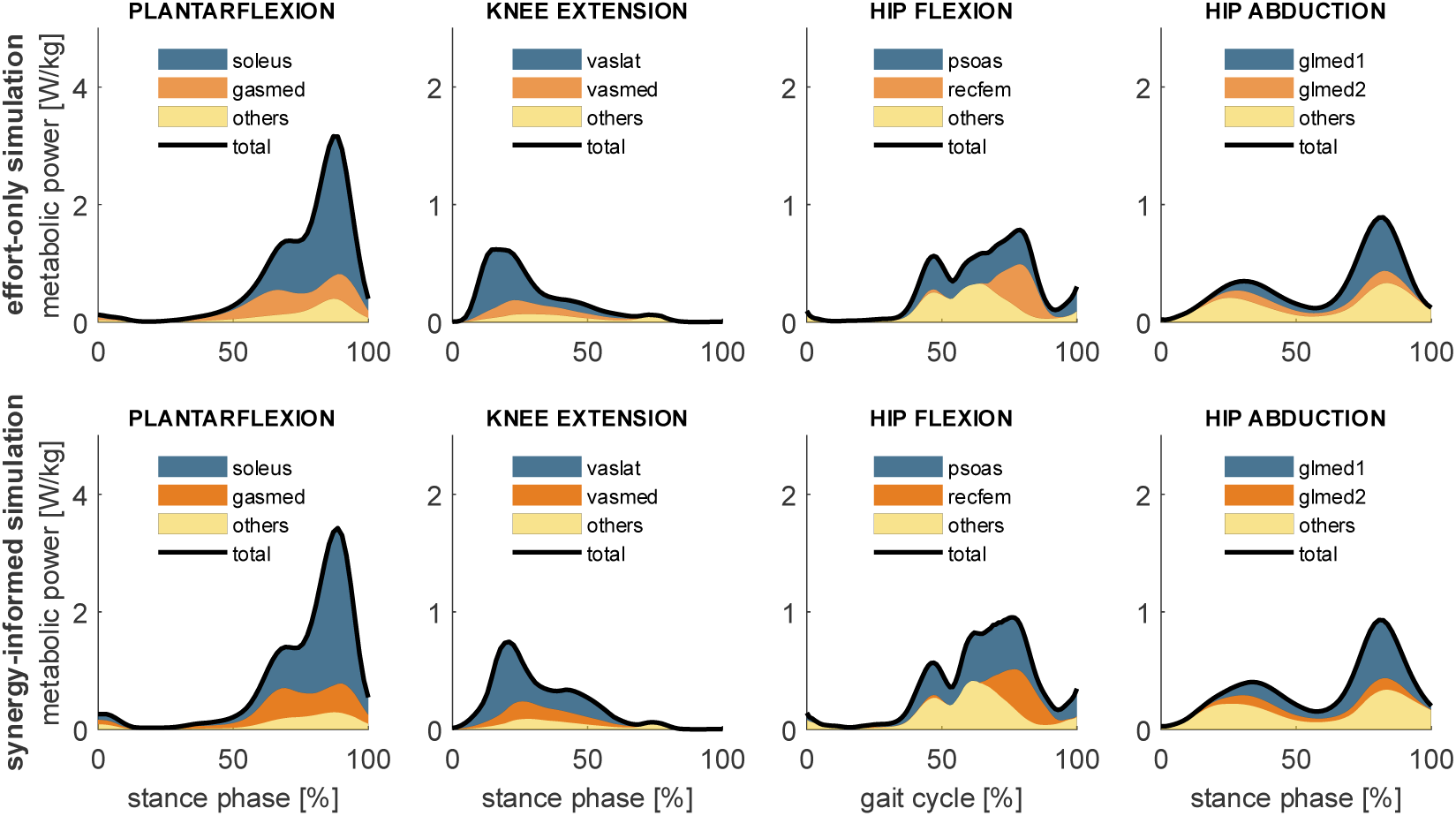
Metabolic power consumed by four muscle groups in unassisted simulations. Results are shown for effort-only simulation (top) and synergy-informed simulation (bottom). Metabolic power (stacked area) highlights the first (slate blue) and second (orange) most expensive muscles; all other muscles were grouped (light yellow). For clarity, values are shown over the stance phase for ankle plantarflexion, knee extension, and hip abduction, and over the entire gait cycle for hip flexion. Mean values were computed across all subjects and gait cycles. Gasmed: gastrocnemius medialis; vaslat and vasmed: vastus lateralis and medialis; recfem: rectus femoris; glmed1 and glmed2: gluteus medius compartments 1 and 2. Gait cycle was defined from heel strike to heel strike of the same limb.

### Synergy weights and activations in the unassisted condition

Four of the six synergies (S1, S2, S4, and S5) had similar timing and activation profiles across subjects (Fig 4). Synergy S1 had a single peak around 50% of the gait cycle (mid-stance), closely following the patterns of the soleus and iliacus. Synergy S2 had a pronounced activation throughout the swing phase, closely following the pattern of the tibialis anterior, with a burst at heel strike. Despite broad consistency, the onset of Synergy S2 was slightly later in subject 2 (∼70% of the gait cycle) than in the other subjects (∼60% of the gait cycle) during the swing phase. Small but non-negligible activations (less than 0.2) were present during the stance phase. Synergy S4 had a single peak around 15% of the gait cycle (early stance). Synergy S5 had pronounced activation at the beginning and end of the gait cycle and a relatively small but consistent peak around 40% of the gait cycle (late stance).

**Fig 4.**
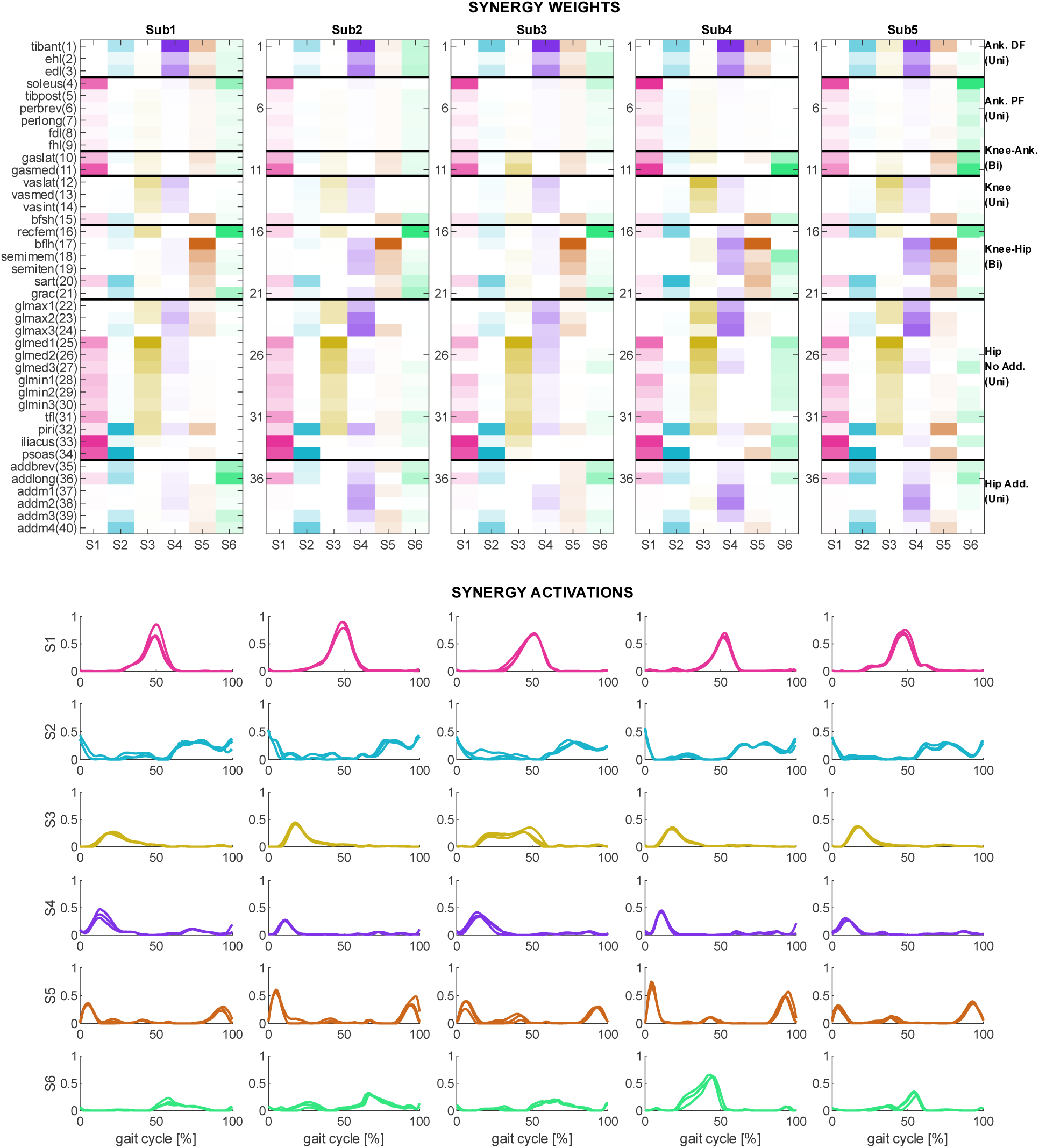
Muscle synergies during unassisted simulations. Time-invariant synergy weights (top) and time-variant synergy activations (bottom) were computed using synergy-informed simulations. Muscle name abbreviations matched those used in the musculoskeletal model (see S1 Appendix (section C)) and were ordered by muscle group (e.g., ankle dorsiflexors) and function (uniarticular or biarticular) to facilitate visualization. Gait cycle was defined from heel strike to heel strike of the same limb. Columns: subjects (e.g., “Sub1” for subject 1); rows: muscles (e.g., “tibant” for tibialis anterior) and synergies (e.g., “S1” for synergy 1).

The remaining two synergies exhibited some subject-specific trends. Synergy S3 was similar in all subjects except one, typically exhibiting a single peak around 20% of the gait cycle (during the loading response), occurring slightly but consistently later than the peak in synergy S4. In subject 3, however, the profile had a longer onset, from the loading response to mid-stance, mimicking the activation of the gluteus medius (Fig B in S2 Appendix). Synergy S6 was similar between subjects 1 and 3, featuring a single peak with long onset and offset times. In subject 2, the pattern was similar, yet it had a small peak around 25% of the gait cycle. In subjects 4 and 5, the pattern was entirely different: subject 4 had a single peak during mid-to-late stance, while subject 5 had two peaks, the one during pre-swing being more pronounced.

Small but visible differences at each synergy activation reflected differences in stride variability. The patterns (specifically, the timing of peaks) were largely preserved between gait cycles. The major differences between gait cycles were the magnitudes, and these differences were synergy specific.

### Optimal torques and metabolic cost reductions

Bilevel optimizations converged within the allocated number of iterations. Bilevel optimization with synergy-informed simulations took 35 min on average, which was nearly 2.3 times longer than effort-only simulations. Both formulations converged to their optima within half of the allocated number of iterations and then did not progress (Fig C in S2 Appendix). Trends in optimal control parameters were joint specific (Figs 5 and 6). Nonetheless, among all subjects and assisted joints, the reduction in average metabolic cost, relative to unassisted conditions, was greater in effort-only simulations than in synergy-informed simulations (Fig 5). In the following subsections, we describe joint-specific trends in the predicted assistive torques and control parameters.

**Fig 5.**
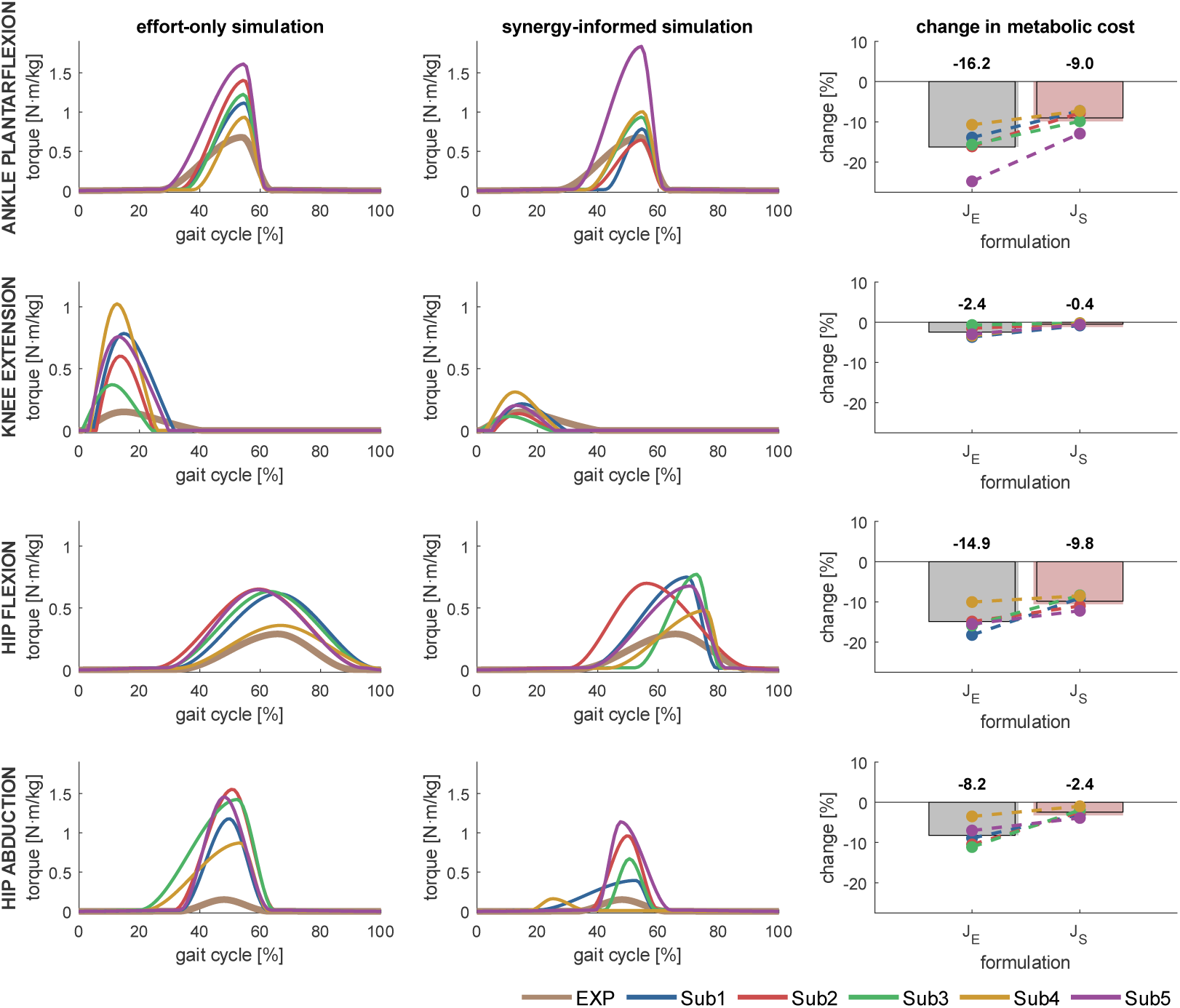
Assistive torques and metabolic cost reductions predicted by effort-only simulations (J_E_) and synergy-informed simulations (J_S_). Torque profiles and average metabolic cost in assisted conditions were computed using bilevel optimization. Changes in metabolic cost were computed relative to unassisted conditions. Optimized torque profiles based on minimal metabolic cost (EXP) were reported from HILO experiments by Poggensee and Collins for ankle plantarflexion assistance [43] and by Franks et al. for knee extension and hip flexion assistance [4], and from a systematic search of optimal peak timing by Park et al. for hip abduction assistance [52]. Gait cycle was defined from heel strike to heel strike of the same limb.

**Fig 6.**
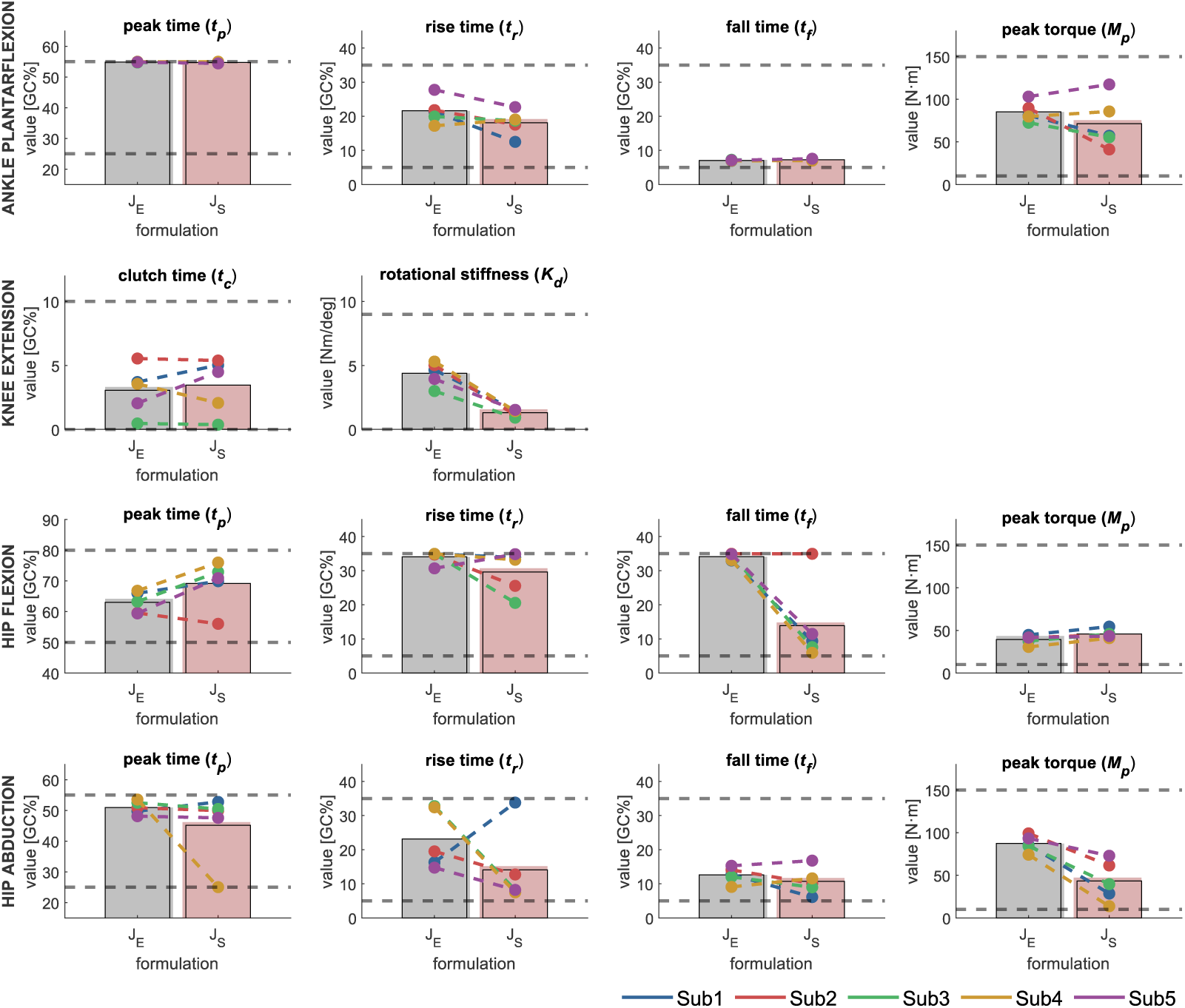
Assistive torque parameters predicted by effort-only simulations (J_E_) and synergy-informed simulations (J_S_). Control parameters were computed using bilevel optimization. Horizontal dashed lines indicate parameter bounds. Gait cycle (GC) was defined from heel strike to heel strike of the same limb.

#### Ankle plantarflexion assistance

Synergy-informed simulations predicted a lower peak torque than the effort-only simulations, resulting in a torque profile similar to HILO-reported values in a subset of subjects. Both formulations predicted optimal torque from mid-stance to pre-swing (Figs 5 and 6). Peak time and fall time were nearly identical between formulations, and rise time was lower in synergy-informed simulations in all subjects except subject 4 (Fig 6 and Table A in S2 Appendix). Peak torque was lower in synergy-informed simulations in subjects 1, 2, and 3 (Fig 6 and Table A in S2 Appendix). In those subjects, assistive torques from synergy-informed simulations better resembled the population-averaged values reported from HILO experiments by Poggensee and Collins [43] than effort-only simulations (Fig 5).

#### Knee extension assistance

Synergy-informed simulations predicted a lower rotational stiffness than the effort-only simulations, resulting in a torque profile similar to HILO-reported profiles in all subjects. The clutch time was subject specific, yet both formulations predicted a timing earlier than 6% of the gait cycle (Fig 6 and Table A in S2 Appendix). The rotational stiffness was lower in the synergy-informed simulations for all subjects, which resulted in a lower peak torque (Fig 6 and Table A in S2 Appendix). The assistive torque predicted by the synergy-informed simulations closely resembled the population-averaged values reported by Franks et al. when assisting the knee joint [4] (Fig 5).

#### Hip flexion assistance

Synergy-informed simulations predicted a peak torque that was slightly but consistently greater than in the effort-only simulations and with more diverse actuation timing. The peak time was greater in the synergy-informed simulations than in the effort-only simulations for all subjects except subject 2; the fall time demonstrated the opposite trend (i.e., the value was lower in the synergy-informed simulations) except in subject 2 (Fig 6 and Table A in S2 Appendix). There was no clear trend in rise time between formulations (Fig 6 and Table A in S2 Appendix). Peak torque in the synergy-informed simulations was slightly greater than in the effort-only simulations for all subjects (Fig 6 and Table A in S2 Appendix). The assistive torque from the effort-only simulations closely resembled the population-averaged actuation timing (i.e., rise time, fall time, and peak time) reported by Franks et al. when assisting the hip joint [4] (Fig 5); however, the peak torque in our simulations was approximately double the experimental peak.

#### Hip abduction assistance

Synergy-informed simulations predicted a lower peak torque than the effort-only simulations, and the peak time was generally consistent between formulations. The peak time was similar between formulations for all subjects except subject 4 (Fig 6 and Table A in S2 Appendix). The rise time in the synergy-informed simulations was lower than in the effort-only simulations for all subjects except subject 1, yet there was a large inter-subject variation (Fig 6 and Table A in S2 Appendix). There was no clear trend in fall time between formulations (Fig 6 and Table A in S2 Appendix). The peak torque in the synergy-informed simulations was lower than in the effort-only simulations for all subjects, yet there was a large inter-subject variation (Fig 6 and Table A in S2 Appendix). The assistive peak time predicted by both formulations closely resembled the population-averaged torque timing reported by Park et al. [52] for all subjects except subject 4 (Fig 5).

### Changes in muscle activations and synergies

Changes in activations were more pronounced in the muscles crossing the assisted joint in both formulations. Yet, the effort-only simulations tended to decrease more muscle activations, relative to unassisted conditions, than the synergy-informed simulations (Fig D in S2 Appendix). Synergy-informed simulations prevented muscle shutdown in only a subset of joints and subjects (Fig 7).

**Fig 7.**
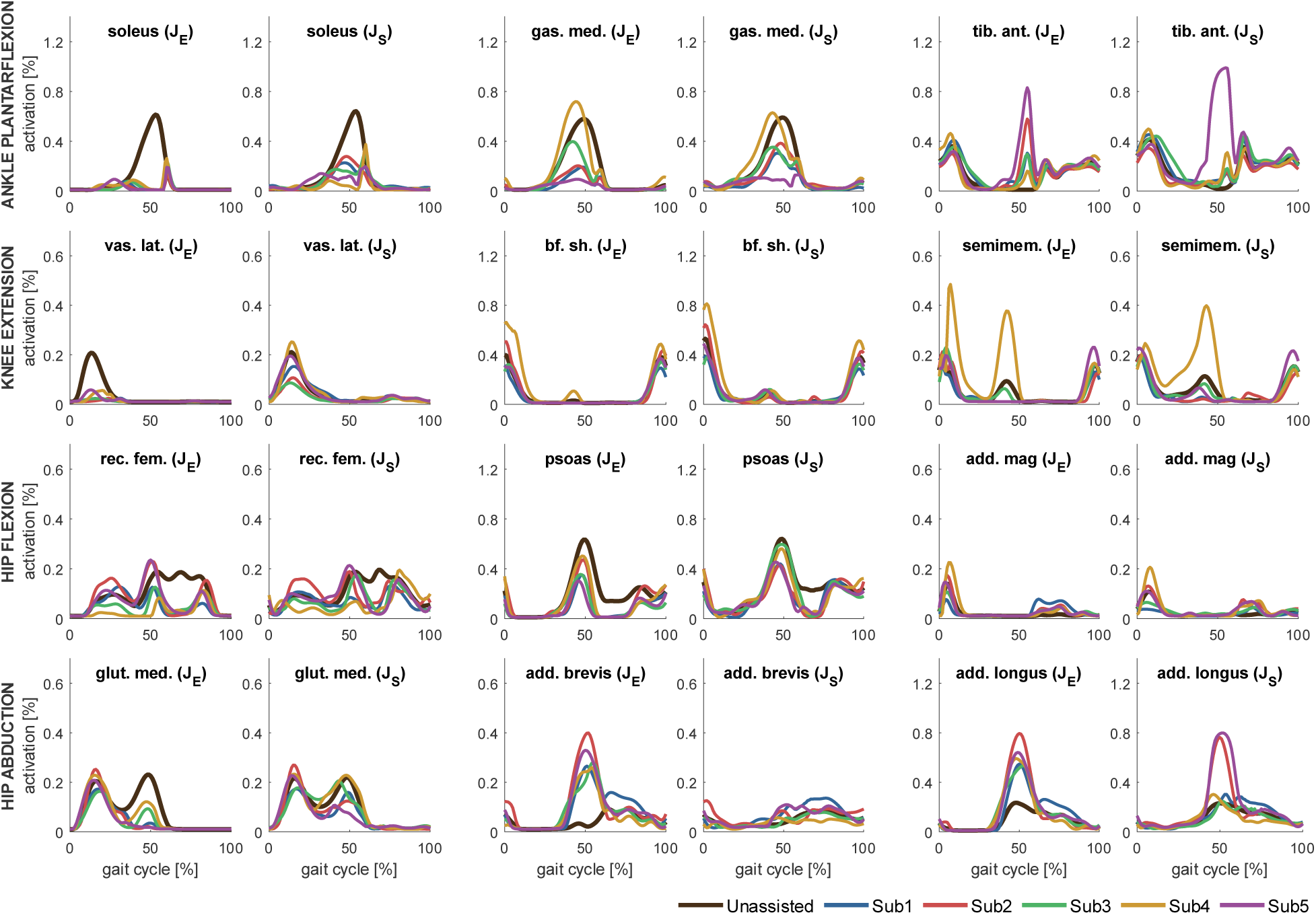
Activations of key muscles in effort-only simulations (J_E_) and synergy-informed simulations (J_S_). These muscles were selected as they either accounted for the greatest metabolic cost during the assisted period or were antagonistic to the assisted joint. Muscle activations were computed using bilevel optimization. Gait cycle was defined from heel strike to heel strike of the same limb.

In the following subsections, we describe changes in muscle activations and synergies, relative to unassisted simulations and between formulations, with emphasis on the muscles that were responsible for the greatest metabolic cost (Fig 3) or that showed the greatest changes in activations (Fig D in S2 Appendix) for each assisted joint.

#### Ankle plantarflexion assistance

Effort-only simulations shut down the soleus during the assisted period whereas synergy-informed simulations did not (Fig 7). In the effort-only simulations, tibialis anterior activation increased relative to the unassisted condition over the assisted period. In the synergy-informed simulations, the soleus activation was not eliminated completely, and the increase in tibialis anterior activation relative to unassisted conditions was less pronounced in 3 of the 5 subjects.

Synergy-mediated coupling prevented substantial reduction of soleus activation in a subset of subjects. In subjects 1, 2, and 3, soleus activation was primarily composed of synergy S1, and gastrocnemius activation was composed of synergies S1, S3, and S5 (Fig 4). As such, decreasing the activation and metabolic cost associated with the ankle plantarflexors involved modulating a broad range of muscles within synergy S1. As a result, only a fraction of soleus and gastrocnemius activations were reduced. On the contrary, in subject 4, soleus activation was composed of synergy S1 and gastrocnemius activation was composed of synergies S1 and S6 (Fig 4), where synergy S6 closely mimicked the gastrocnemius activation profile. As a result, in order to reduce soleus activity (the primary driver of metabolic cost), the optimizer substantially decreased synergy S1 activity and increased synergy S6 activity (Fig 8). This compensation strategy enabled a substantial reduction of soleus activation and an increase of gastrocnemius activation (Fig 7). In subject 5, soleus activation was composed of synergies S1 and S6, where synergy S6 contributed to soleus and hip abductor muscles (Fig 4). Thus, the optimizer substantially reduced synergy S1 activity and suppressed synergy S6 activity to nearly zero. This compensation strategy enabled a substantial reduction in the activity of all plantarflexor muscles and also decreased gluteus medius and minimus, an effect that was not observed in the effort-only simulations (Fig D in S2 Appendix and Table B in S2 Appendix). In subject 5, synergy S2, which strongly drove the tibialis anterior, was upregulated as a compensatory response toward near-maximal levels during push-off, to a slightly greater magnitude in the synergy-informed simulations than in the effort-only simulations (Fig 7, Table B in S2 Appendix).

**Fig 8.**
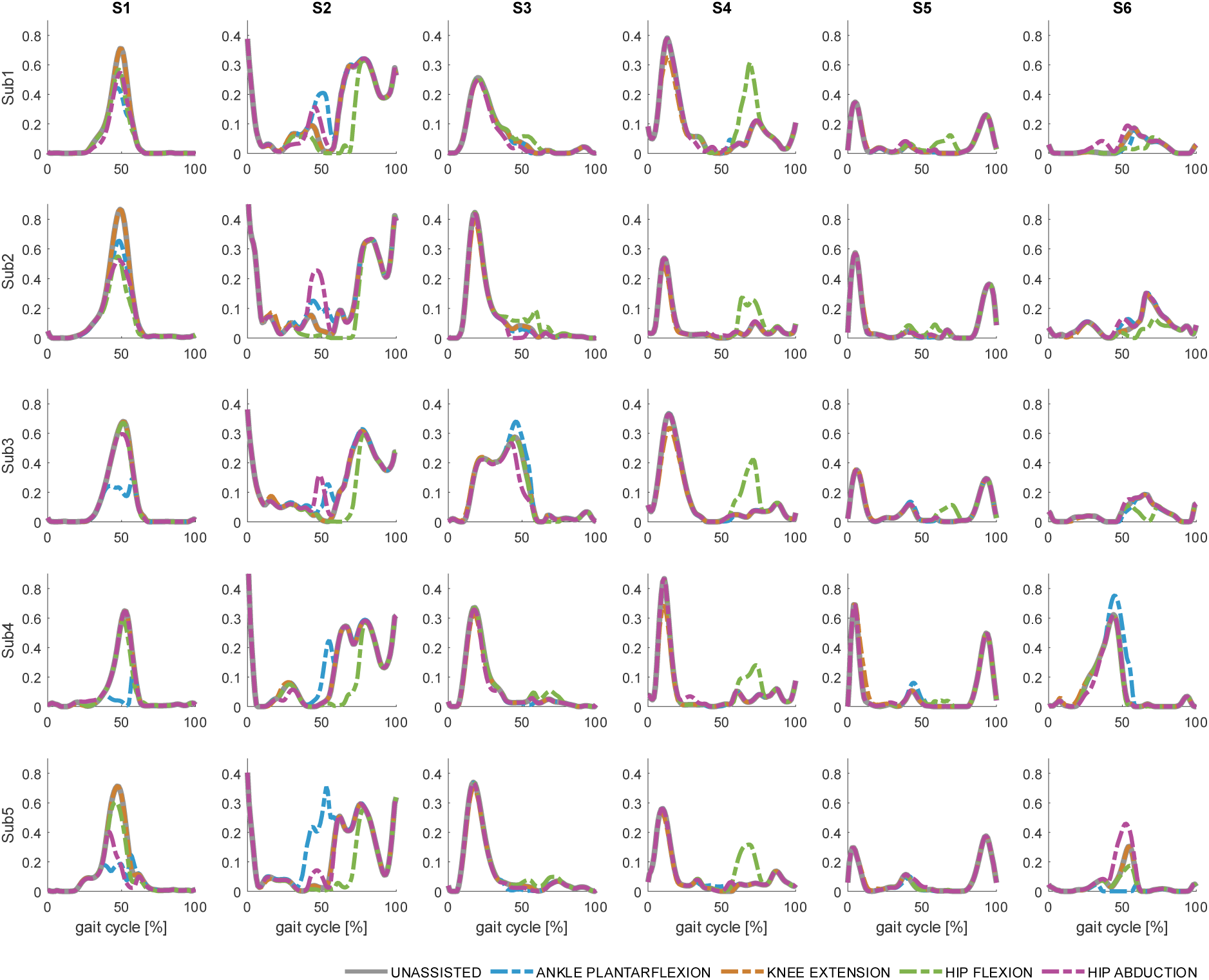
Synergy activations during simulations of unassisted and assisted gait. Synergy activations were computed using bilevel optimization with an inner loop based on synergy-informed simulations. Gait cycle was defined from heel strike to heel strike of the same limb. Columns: synergies (e.g., “S1” for synergy 1); rows: subjects (e.g., “Sub1” for subject 1).

#### Knee extension assistance

Effort-only simulations shut down the vastus muscles during the assistive period whereas synergy-informed simulations did not (Fig 7). In effort-only simulations, a small increase in antagonist muscle activations was observed relative to unassisted conditions (e.g., ∼7% increase in biceps femoris short head activation). In the synergy-informed simulations, the increase in antagonist muscle activations was less pronounced (e.g., ∼2% increase in biceps femoris short head activation).

The synergy-mediated coupling prevented a substantial reduction in vastus activations for all subjects. These muscles were composed of synergies S3 and S4 (Figs 4 and 8). Such a broad range of synergy contributions discouraged the optimizer from suppressing the vastus lateralis, the primary driver of metabolic cost. Overall, the synergy-informed simulations resulted in relatively small changes in muscle activations and across fewer muscles compared to the effort-only simulations (Fig D in S2 Appendix and Table C in S2 Appendix).

#### Hip flexion assistance

Both formulations predicted nearly zero agonistic muscle activations during the assisted period (Fig 7). Changes in muscle activations, relative to the unassisted simulations, were more pronounced in the effort-only simulations than in the synergy-informed simulations (Table D in S2 Appendix). Both formulations primarily decreased rectus femoris, iliacus, psoas, and adductor brevis and longus activations (Fig 7 and Fig D in S2 Appendix). In the synergy-informed simulations, a small decrease in ankle plantarflexor activations (excluding soleus) was observed (e.g., ∼13% decrease in tibialis posterior activation); in the effort-only simulations, the opposite trend was observed (e.g., ∼4% increase in tibialis posterior activation).

The synergy-mediated coupling did not prevent a substantial reduction in rectus femoris activation during the assisted period in all subjects. Synergy S2 substantially contributed to the rectus femoris peak, which occurred during early swing. The optimizer decreased this synergy to zero during the assisted period while increasing others (primarily synergy S4) as a compensatory response (Fig 8).

#### Hip abduction assistance

The effort-only simulations substantially reduced gluteus medius activation during the assistive period whereas the synergy-informed simulations did not (Fig 7). Similar to hip flexion assistance, the effort-only simulations predicted more pronounced changes in muscle activations, relative to the unassisted condition, than the synergy-informed simulations (Fig D in S2 Appendix and Table E in S2 Appendix). Both formulations primarily decreased gluteus and iliacus activations and increased those of adductors longus, brevis, and magnus (Fig 7 and Fig D in S2 Appendix). Yet, in the synergy-informed simulations, the increase in antagonistic muscle activations varied substantially across subjects and muscles. For instance, in 2 of the 5 subjects, adductor longus activations increased by ∼40%, similar to the effort-only simulations, while in the other subjects this increase was only ∼10%. In contrast, the adductor brevis activation increased by only ∼10% and was relatively consistent among subjects.

An increase in hip adductor activations was a compensatory strategy to generate hip flexion torques. Gluteus medius was the primary driver of metabolic cost in the hip abductor muscle group (Fig 3). Decreasing metabolic cost led to a decrease in the activations of gluteus medius and other hip abductors such as iliacus, which generated both hip abduction and hip flexion moments. Therefore, offloading iliacus required other muscles to generate greater hip flexion moments. This compensatory strategy was required since, in inverse simulations, muscle moments must reproduce the net joint torques computed by inverse dynamics (as in the unassisted conditions). In the effort-only simulations, the compensatory strategy involved increasing the adductor muscle activations, primarily adductors brevis and longus, to generate hip flexion torques during terminal stance (Fig E in S2 Appendix).

Synergy-mediated coupling prevented the optimizer from substantially reducing the gluteus medius activation in all subjects. This muscle was prominently represented by synergy S1. Thus, the optimizer decreased synergy S1, which coupled several hip abduction and ankle plantarflexion muscles. In subjects 1, 3, and 4, the reduction of synergy S1 was relatively small (less than 0.1) as the synergy-mediated coupling involved the modulation of several muscles, primarily across the hip abductors. In subjects 2 and 5, however, the reduction of synergy S1 was substantially greater (0.25 and 0.5, respectively) and resulted in a substantial increase in adductor longus activation, similar to the effort-only simulations (Fig 7 and Table E in S2 Appendix). In these two subjects, the coupling mediated by synergies did not substantially spread the modulation of activations; hence, the use of the adductor longus to generate hip flexion torques, as in the effort-only simulations, was also observed (Fig E in S2 Appendix).

## Discussion

In this study, we examined whether synergy-informed simulation would prevent non-physiological muscle shutdown during assistance and predict torque profiles that were consistent with experimental values. Our results revealed a joint-specific effect governed by synergy weights. With ankle plantarflexion or knee extension assistance, effort-only simulations shut down agonist muscle activations during the assisted period, which was not observed in synergy-informed simulations. In 3 of the 5 subjects, peak ankle plantarflexion torques in the synergy-informed simulations (0.79 N·m/kg, mean) were lower than those in the effort-only simulations (1.25 N·m/kg, mean) and were more consistent with HILO experiments (0.68 N·m/kg, mean) [43]. In all subjects, peak knee extension torques in the synergy-informed simulations (0.19 N·m/kg, mean) were lower than in the effort-only simulations (0.71 N·m/kg, mean) and were more consistent with HILO experiments (0.15 N·m/kg, mean) [4]. With hip flexion assistance, both formulations predicted a substantial reduction of agonist muscle activations and overestimated the peak assistive torque (0.58 N·m/kg in effort-only simulations and 0.68 N·m/kg in synergy-informed simulations) in comparison to HILO experiments (0.29 N·m/kg, mean) [4]. With hip abduction assistance, the effort-only simulations substantially reduced agonist (gluteus medius) activation, which was not observed in the synergy-informed simulations. However, 2 of the 5 subjects showed substantial recruitment in some antagonist muscles. No experimental study has optimized peak torque for hip abduction assistance based on minimal metabolic cost, though the peak assistive times predicted in the present study were consistent between formulations and reported experimental values (∼50% of the gait cycle) [52]. These findings have direct implications for determining the situations in which simulation-based approaches should be used to inform wearable robot design and where their predictions may be unreliable.

### Synergy structure: Modulation of muscle activations and metabolic cost

The extracted synergy structure was partially consistent with motor modules reported in EMG-based decompositions of human walking. Six synergies were required to achieve a VAF of 95%, on average, which was within the range that is typically reported for walking (4–6 modules) [20,21,45]. Four of the six synergies were consistent across subjects in timing and profile, capturing the principal activation events of the gait cycle: mid-stance activation (synergy S1), a swing-phase activation (synergy S2), an initial contact activation (synergy S4), and a double-peak activation at the beginning and end of the gait cycle (synergy S5). These timings aligned with synergy activation patterns extracted from EMG recordings [20,21] although some discrepancies remained. For instance, we did not observe a synergy whose activation peaked during early swing (∼62–75% of the gait cycle) [20,21]. One possible explanation is that synergies extracted from a single-task, minimal-effort baseline may not fully capture the coordination strategies present during natural walking. Notably, a synergy with an early-swing peak was also absent from motor primitives extracted under exoskeleton assistance [24], suggesting that the discrepancy is not unique to our baseline. This finding highlights the sensitivity of the extracted synergies to the decomposition methodology, which makes it difficult to assess whether our synergy modules were physiologically grounded without confounding differences in the number of analyzed muscles and extraction methods [54–56].

Development of methods for robust extraction of synergies remains a promising direction for capturing physiologically plausible motor coordination. Current NMF-based approaches are mathematically constrained to identify no more synergies than the number of analyzed features. This approach may cause the extracted synergy set to underrepresent the coordination repertoire truly available to an individual [57]. For instance, according to the synergy expansion hypothesis, different individuals may develop distinct synergy solutions for a given motor task, depending on their motor history, implying that synergy weights extracted from a single-task baseline may reflect only a subset of the available motor repertoire. Whether extraction from richer and more diverse behavioral contexts (or, on the contrary, extraction of synergies in higher domains [58]) would yield a more generalizable and complete representation of neuromuscular organization remains an open question and a promising direction for future investigation.

Overestimation of metabolic cost in synergy-informed simulations might reflect muscle model limitations rather than an implausible synergy structure. Informing muscle control with synergy modules increased co-activation, consistent with prior modeling predictions [29,30,45]. Hence, metabolic cost was also expected to be greater in synergy-informed simulations than in effort-only simulations and was expected to increase with decreasing synergy dimension. Yet, the magnitude of the overestimation (32% with six synergies in the unassisted condition) did not necessarily stem only from increases in co-activation. Musculoskeletal models based on Hill-type muscle actuators are known to overestimate mechanical power due to simplified representations of muscle–tendon excursion [38,59] and muscle positive mechanical power is directly proportional to metabolic cost [49,60]. As such, metabolic overestimation was consistent with an intrinsic limitation of the muscle–tendon actuator model and was not necessarily evidence that the synergy modules yielded muscle activations that were less physiologically plausible than in the effort-only simulations.

### Synergy coupling discourages muscle shutdown with assistance and plays a key role in determining optimal assistance

As hypothesized, when muscles shared synergy weights, reducing the activation of any single muscle required modulating the activity of all muscles coupled within the same synergy. This constraint prevented the optimizer from suppressing individual muscles in isolation, partially offsetting the unrealistic metabolic benefit predicted with the effort-only formulation and discouraging unrealistically large assistive torques. Notably, a high degree of load sharing can be achieved by increasing the exponent in the effort-minimization objective function [10,61], which suggests another modeling choice to attenuate peak assistive torques. However, a high degree of load sharing degrades agreement between predicted muscle activations and EMG in unassisted conditions [62,63]. In this regard, solving the muscle redundancy problem based on minimal muscle activations with a low degree of load sharing and constrained by synergy modules exhibited a valuable emergent property: simulated muscle activations preserved agreement with EMG during unassisted gait yet imposed implicit load sharing during assisted gait through synergy constraints. This property held only when the synergy weight prevented individual muscle isolation, i.e., when a reduction of the activation in the target muscle group (e.g., plantarflexion assistance) required modulating synergies that also governed other metabolically expensive muscles (e.g., soleus and gastrocnemius).

The synergy weights directed the modulation of muscle activations and the prediction of peak assistive torque. For motor-like actuation, the value of the peak time parameter was determined by the instant at which metabolic power was greatest, which was common between effort-only and synergy-informed simulations (Fig 2). In contrast, the value of the peak torque parameter was determined by synergy-mediated muscle coupling: a structural barrier against individual muscle suppression was created when metabolically expensive muscles were coupled with many other muscles. To investigate this effect further, we conducted a post-hoc analysis optimizing ankle plantarflexion and hip abduction assistance for subject 5 using synergy weights from subject 1. In the original synergy weights of subject 5, the optimizer substantially induced antagonist muscle recruitment. When synergy weights of subject 1 were applied instead, this compensation was suppressed: peak assistive torques and antagonist activations were lower in both ankle plantarflexion and hip abduction (Fig 9), confirming the influence of the synergy weights on the predicted peak assistive torque.

**Fig 9.**
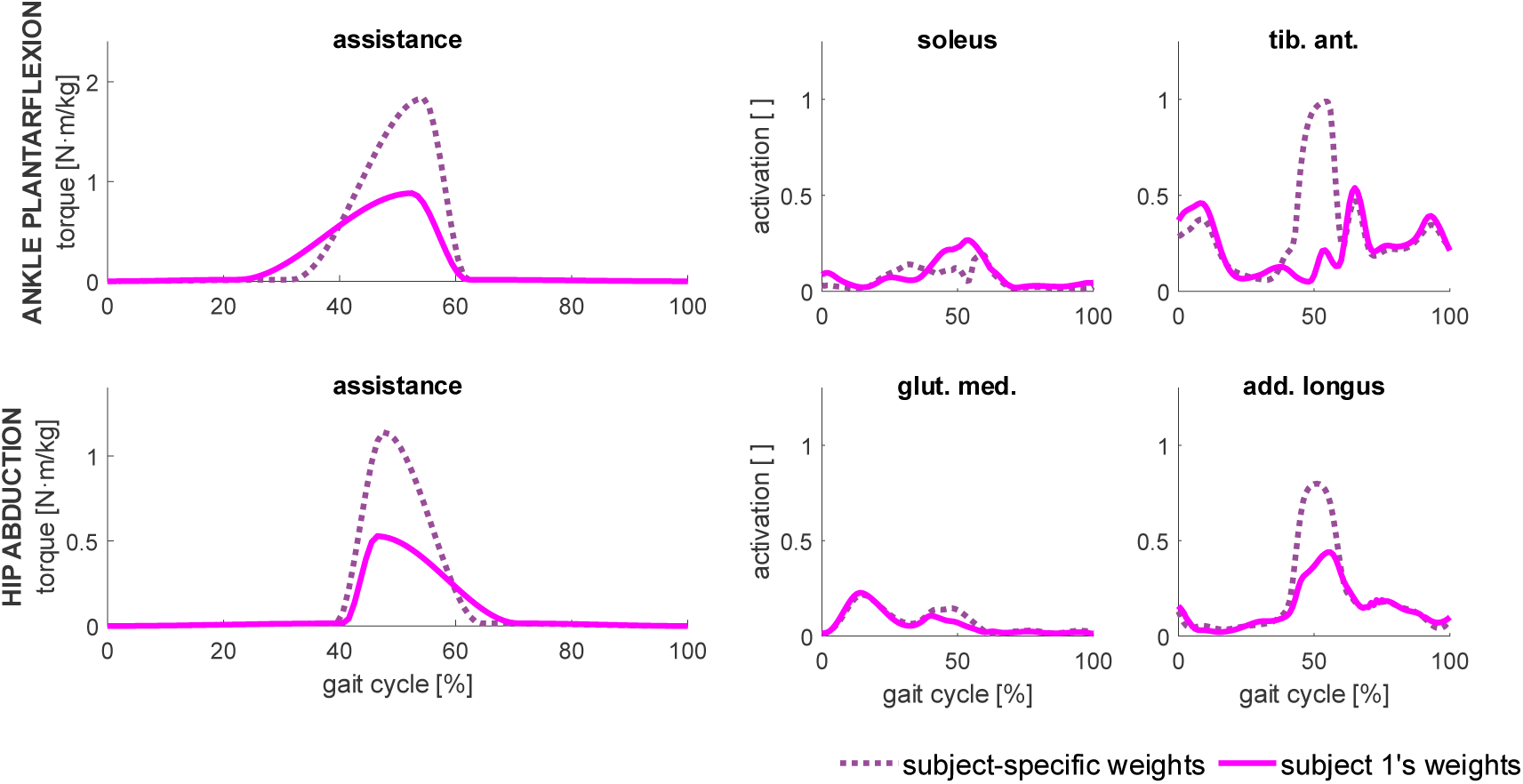
Optimal assistance using two sets of synergy weights. Optimal torques (left) and muscle activations (right) during gait with ankle plantarflexion assistance (top) and hip abduction assistance (bottom) in subject 5, using the synergy weights derived from subject 5’s unassisted simulations (“subject-specific weights,” dotted lines) and from subject 1’s unassisted simulations (“subject 1’s weights,” solid lines). Gait cycle was defined from heel strike to heel strike of the same limb.

Recent studies that developed exoskeleton control based on synergy-informed muscle control have shown promising results in real-world deployment [25,64]. Our results support the use of synergy-based approaches to inform assistive strategies. Yet, we highlight the importance of ensuring that the synergy modules represent realistic muscle coordination. In particular, we suggest examining whether the assisted muscle group is driven by a single synergy or several, and whether those synergies also drive other metabolically expensive muscles. When a metabolically expensive muscle is governed by a single, otherwise uncoupled synergy, its shutdown remains the optimal solution for minimizing metabolic cost and the synergy constraint offers little protection against predicting unrealistically high assistive peak torques.

### Plausibility of synergies as the only missing motor control property

Synergy-informed simulations captured the trends observed in experimental muscle coordination patterns during optimal assistance, yet important discrepancies remained. No study has examined muscle synergy structures under HILO-derived optimal assistance, which limited the degree to which our predictions could be validated. Poggensee and Collins [65] have provided the most detailed characterization of muscle activity under optimal ankle plantarflexion assistance, and their findings are partially consistent with our predictions. They reported an approximately 50% reduction in soleus activation relative to their “zero torque” condition (i.e., with the device worn but unpowered), which agreed with the reduction in soleus activation that we predicted in subjects where synergy coupling prevented muscle shutdown. Poggensee and Collins also showed that changes in muscle coordination were not limited to muscles crossing the ankle, which was consistent with another study that optimized assistance across multiple joints [4]. This muscle adaptation strategy was consistent with the prediction that synergy coupling would distribute coordination changes broadly across the leg. However, a key discrepancy remained: Poggensee and Collins reported no increase in tibialis anterior activation during push-off, whereas our simulations predicted this increase as the primary mechanism that limited the peak assistive torque. This discrepancy suggests that synergy coupling may not be the only neuromuscular component missing from effort-based simulations to improve the accuracy of predicted human–robot interaction.

The remaining discrepancies may reflect neuromuscular control pathways that are missing from effort-minimization frameworks. Muscle coordination during locomotion is thought to emerge from the combined action of spinal central pattern generators (CPGs), which provide rhythmic feedforward motor commands [66,67] and peripheral sensory feedback loops that continuously modulate the timing and amplitude of muscle recruitment throughout the gait cycle [68]. Effort-minimization simulations captured neither mechanism directly: they generated an efficient solution without the sensorimotor feedback that shaped how the nervous system adapted coordination. Explicitly modeling feedback and feedforward control pathways may be the most mechanistically complete approach to predicting muscle dynamics under assistance, yet there are several practical limitations. For example, reflex-based and CPG-based models require forward simulations that are orders of magnitude more computationally expensive than inverse approaches [32,69]. The models also contain numerous parameters (e.g., reflex gains, sensory delays, and threshold functions) that cannot be directly measured and require indirect parameter identification [70]. Furthermore, reflex gains that are calibrated during unassisted gait may not generalize to assisted conditions where afferent signals are substantially altered. In sum, modeling more accurate muscle control behavior remains a fruitful field of study, and muscle synergies represent a practical approach for predicting more realistic assistive strategies than other approaches, despite being an incomplete representation of human motor control.

### Limitations and future work

This study has four major limitations. First, the sample size of five subjects limits the generalizability of some observations. The main finding concerned the introduction of synergy-mediated coupling, where the distribution of muscle activations across synergies prevented the optimizer from suppressing the activity of individual muscles during assisted gait. A larger cohort of subjects would not affect this conclusion but would provide more robust estimates of how frequently and to what extent the synergy modules prevented muscle shutdown across assistive devices, joints, and muscles. Second, the inverse simulation approach prescribed kinematics from unassisted gait, which was a deliberate choice to compare effort-only and synergy-informed muscle control in isolation, at the expense of ignoring kinematic adaptation to assistance. Predictive simulations represent a state-of-the-art approach to predicting adaptation to exoskeleton assistance. However, several studies have shown that predicted changes in kinematics were substantially greater than those observed experimentally [71,72], which suggests that critical physiological constraints have not been included in these simulations. Future predictive simulations may benefit from incorporating synergy-informed muscle control, following the considerations described in the Discussion (synergy coupling), to generate more realistic movement patterns and avoid isolated suppression of individual muscles during assisted gait. Third, all devices in this study were modeled as ideal torque generators, neglecting actuator mass, inertia, and bandwidth. Introducing actuator dynamics is a promising direction that may narrow the gap between predicted optimal torques and observations from HILO experiments. Finally, although the musculoskeletal model was scaled to match the anthropometry of the subjects [34] and the musculotendon parameters were calibrated to ensure agreement between simulated muscle excitations and EMG recordings during gait [38], the models used in this study (including their muscle paths, muscle–tendon dynamics, and metabolics) are nevertheless only approximations of reality. Repeating the present study using different musculoskeletal, muscle–tendon, muscle activation, and metabolic models would further test the robustness of this study’s findings.

### Conclusions

This study indicates that the physiological plausibility of simulation-optimized exoskeleton assistance was joint specific, determined by how the muscles at each assisted joint were coupled within synergies. When the synergy governing a metabolically expensive muscle also governed other expensive muscles or spread its activation across many muscles (as we observed during ankle plantarflexion, knee extension, and hip abduction assistance), the synergy-informed formulation prevented suppression of single muscles and predicted torque profiles that were more consistent with human-in-the-loop experiments than the effort-only formulation. When this coupling was absent, as we observed during hip flexion assistance, the synergy constraint offered little protection and peak torque predictions remained unrealistically high. These findings establish synergy weights as a potential predictor of simulation accuracy and provide guidance for when synergy-informed simulations may be useful for assistive device design. However, a synergy-informed motor control structure does not fully provide the physiological constraints that are missing from pure effort-minimization approaches. As simulation-driven approaches become increasingly central to systematic wearable robot development, establishing joint-specific trust boundaries for simulation tools is a necessary foundation for scalable, reliable, and translatable device design.

## Supporting information

**S1 Appendix. Supplementary methods.** (A) Synergy constraint formulation, comparison with existing approaches, and selection of the synergy deviation weight. (B) Control parameter bounds and temporal constraints. (C) Names and abbreviations of the muscles in the musculoskeletal model.

**S2 Appendix. Supplementary figures and tables.** Fig A, initial-guess sampling; Fig B, muscle activations across formulations and synergy dimensions; Fig C, computation time and optimizer iterations; Fig D, changes in average muscle activations; Fig E, hip joint moments during hip abduction assistance. Table A, optimal control parameters per subject; Tables B–E, changes in muscle activations for each assisted joint.

